# Spatial Transcriptomics Arena (STAr): an Integrated Platform for Spatial Transcriptomics Methodology Research

**DOI:** 10.1101/2023.03.10.532127

**Authors:** Xi Jiang, Danni Luo, Esteban Fernández, Jie Yang, Huimin Li, Kevin W. Jin, Yuanchun Zhan, Bo Yao, Suhana Bedi, Guanghua Xiao, Xiaowei Zhan, Qiwei Li, Yang Xie

## Abstract

The emerging field of spatially resolved transcriptomics (SRT) has revolutionized biomedical research. SRT quantifies expression levels at different spatial locations, providing a new and powerful tool to interrogate novel biological insights. An essential question in the analysis of SRT data is to identify spatially variable (SV) genes; the expression levels of such genes have spatial variation across different tissues. SV genes usually play an important role in underlying biological mechanisms and tissue heterogeneity. Currently, several computational methods have been developed to detect such genes; however, there is a lack of unbiased assessment of these approaches to guide researchers in selecting the appropriate methods for their specific biomedical applications. In addition, it is difficult for researchers to implement different existing methods for either biological study or methodology development.

Furthermore, currently available public SRT datasets are scattered across different websites and preprocessed in different ways, posing additional obstacles for quantitative researchers developing computational methods for SRT data analysis. To address these challenges, we designed Spatial Transcriptomics Arena (STAr), an open platform comprising 193 curated datasets from seven technologies, seven statistical methods, and analysis results. This resource allows users to retrieve high-quality datasets, apply or develop spatial gene detection methods, as well as browse and compare spatial gene analysis results. It also enables researchers to comprehensively evaluate SRT methodology research in both simulated and real datasets. Altogether, STAr is an integrated research resource intended to promote reproducible research and accelerate rigorous methodology development, which can eventually lead to an improved understanding of biological processes and diseases. STAr can be accessed at https://lce.biohpc.swmed.edu/star/.

## 1 Introduction

Spatially resolved transcriptomics (SRT) has revolutionized genomics research. It enables high-throughput quantitative assessment of the location and abundance of gene activity within a tissue, which can elucidate molecular mechanisms at an unprecedented level of spatial detail. Traditional molecular profiling technologies (e.g., single-cell or single-nuclei RNA sequencing) dissociate tissues, lose the spatial context of gene expression [1–3], or offer limited gene-level assessment via tissue slides. In contrast, SRT technologies empower the comprehensive characterization of molecular abundance within individual cells while preserving spatial information. SRT technologies are increasingly used to discover location-specific gene function, facilitating novel biological discoveries and disease studies [4–8]. The rapid adaptation of SRT had led to large volumes of data, which can potentially lead to further novel biological insights; however, several limitations prevent the research community from fully utilizing SRT technologies and data resources. These limitations include time spent retrieving data from different sources, data preprocessing and quality control steps, evaluation and selection of proper data analysis methods, and interactive exploration of analysis results across studies and methods. To overcome these challenges, we created a unique resource that we call Spatially Resolved Transcriptomics Arena (STAr), an integrated and user-friendly platform for data collection, analysis, and visualization.

We extensively collected SRT data from seven technologies: 1) Sequential fluorescence *in situ* hybridization (FISH) (seqFISH/seqFISH+) [9], an imaging-based technology that enables the identification of many RNA transcripts sequential rounds of hybridization; 2) Multiplexed error-robust FISH (MERFISH)[10], another FISH technology with fewer fluorescent channels and more hybridization rounds compared to seqFISH; 3) Spatial transcriptomics (ST) [11] and 4) high-definition spatial transcriptomics (HDST) [12], two recently developed SRT approaches that obtain high-quality RNA-sequencing data with positional information through arrayed probes with unique positional barcodes. 10x Genomics, a biotechnology company, has developed a new molecular profiling solution called the Visium Spatial Gene Expression, which provides a growing number of downloadable spatial transcriptomics data and a platform for users to perform spatial transcriptomics; 5) STARmap (spatially-resolved transcript amplicon readout mapping) [13], a 3D intact-tissue RNA sequencing approach at single-cell resolution; 6) Slide-seqV2 [14], an improved version of Slide-seq, which has a higher RNA capture efficiency; and 7) An integration of a microarray-based SRT method and a single-cell RNA sequencing method (denoted as Other) [15], a recently developed method to characterize the cell populations and their spatial organization in a tissue. The development of these SRT techniques has provided biological insights for many fields, including neuroscience [16], disease pathology [1], and embryonic development [17].

In addition, we extensively surveyed important SRT analysis methods with a focus on spatially variable (SV) gene detection. These methods belong to four classes: 1) Gaussian process-based approaches, including SpatialDE [18], SPARK [19] and BOOST-GP [20], which model the spatial correlation among gene expression levels via various kernel functions in the Gaussian process; 2) Methods based on marked point processes, including Trendsceek [21]; 3) Energy-based methods, represented by BOOST-MI [22], which uses Hamiltonian energy to characterize spatial patterns in the Ising model; and 4) Non-model-based methods, such as BinSpect [23], which is based on statistical enrichment of spatial network neighbors. Another non-model-based method, SpaGCN, [24] is based on a graph convolutional network and domain-guided differential analysis. These SV gene identification approaches facilitate the comprehensive exploration of SRT datasets and contribute discoveries to further genomics research. To provide an unbiased and comprehensive assessment of the performance of different SV gene identification methods, we have applied each SV gene identification method to all real datasets collected here, in addition to a large number of synthetic datasets generated under different scenarios (e.g., different spatial patents and percentages of zero counts). These results can be utilized to guide the selection of appropriate analysis methods for specific applications, as well as facilitate methodology development.

Furthermore, we have created an open platform containing state-of-the-art datasets, methods, and analysis results. We present a website (https://lce.biohpc.swmed.edu/star/) comprising three main functions: 1) A collection of organized and pre-processed SRT datasets including both simulated and real datasets; 2) A comprehensive comparison of existing methods for SV gene identification with reproducibility and documentation; and 3) A user-friendly interface of spatial analysis results and visualization of real SRT datasets. Moreover, this open resource allows users to apply their proposed methods to the rich datasets provided on the platform and compare performance with other existing methods.

Curated across a broad scope, our SRT resources are readily available to the wider research community to facilitate data retrieval, selection of the most suitable spatial analysis methods, and exploration of biological results without requiring specific programming experience or knowledge. Importantly, this platform is extensible, allowing researchers from data scientists to biologists to contribute expertise. By allowing the external contribution of datasets, methods, and analysis results, STAr is an open data platform that facilitates long-term spatial molecular research.

## 2 Results

### 2.1 An Overview

STAr is a resource for spatially resolved transcriptomics research, and its workflow is depicted in Figure 1. We provide three components: 1) Curated datasets from real studies or simulations; 2) Reproducible statistical analysis methods; and 3) Interactive, explorable analysis results. STAr is intended to be an open platform extending beyond our local effort. As such, we also provide contribution guidelines for the wider research community for the inclusion and integration of additional spatial transcriptomics datasets, models, and analysis results.

**Fig. 1.**
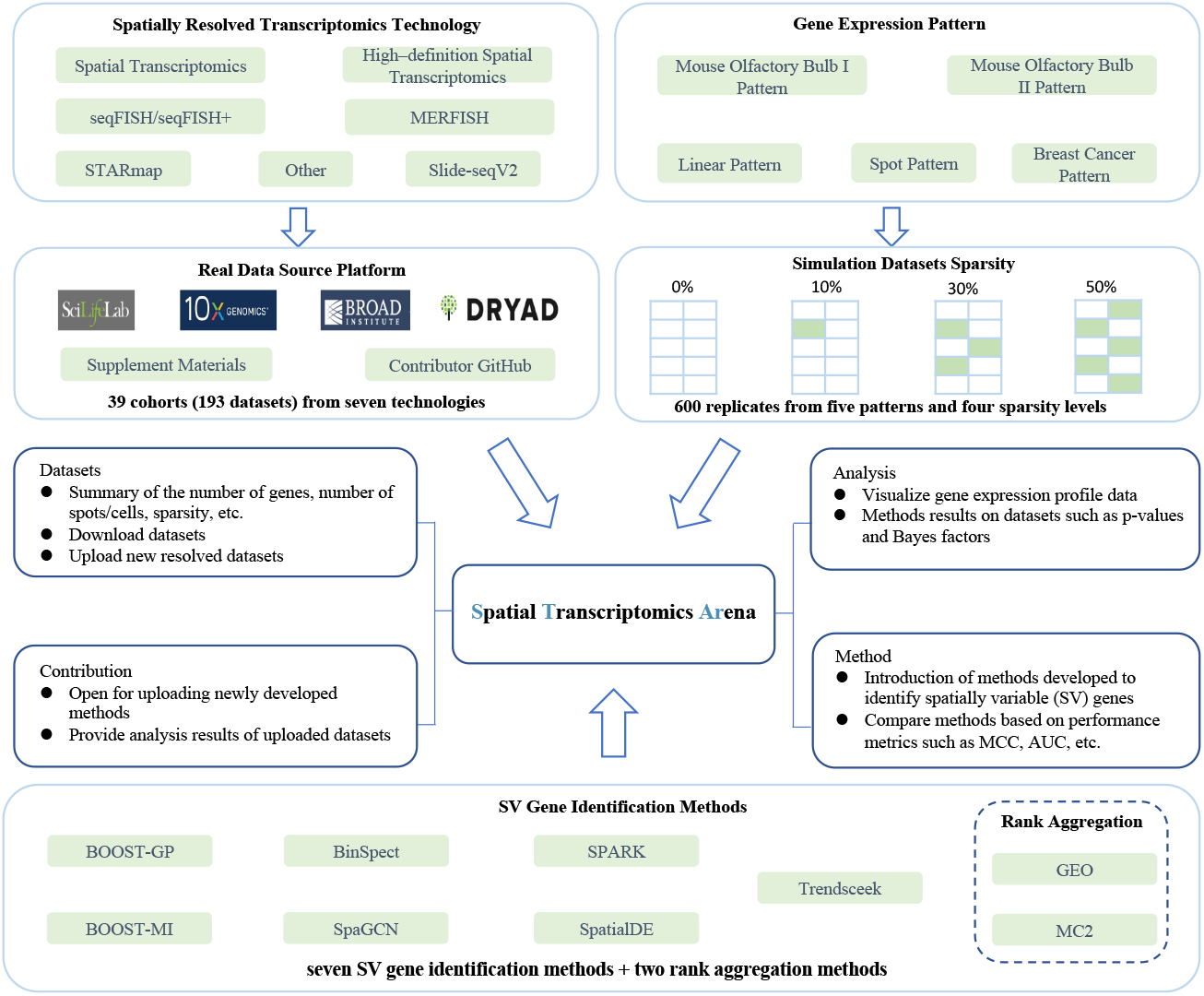
The STAr (Spatial Transcriptomics Arena) workflow: 1) We collected real and simulated datasets. For real datasets, we retrieved publicly available datasets from seven spatially resolved transcriptomics technologies. Simulated datasets were generated using the settings of five spatial gene expression patterns with four levels of sparsity. 2) We surveyed seven existing spatial gene analysis methods and provided two rank aggregation methods. 3) We performed analysis for combining datasets and methods, and the analysis results are tabulated and visualized.

### 2.2 A Curated Collection of SRT Datasets

We collected publicly available SRT datasets directly from SciLifeLab (https://www.scilifelab.se/data), 10x Genomics (https://www.10xgenomics.com/products/spatial-gene-expression), the Broad Institute of MIT and Harvard (https://www.broadinstitute.org/), Dryad (https://datadryad.org), and GitHub repositories. These datasets are resolved by two types of SRT technologies: sequencing-based technologies, such as ST [11], its improved 10x Visium platform, and HDST [12], and imaging-based technologies, such as seqFISH [9], seqFISH+ [25], and MERFISH [26].

In total, we curated 193 datasets (including replicates, layers, samples or sections) across 39 cohorts, which are available for download on our website, with a detailed summary in Table S1 within the Supplementary Notes. Resolved datasets cover two organisms: the human and mouse, encompass 16 tissues from the brain, spinal cord, and ovaries, and eight diseases such as heart failure, melanoma, human breast cancer, etc. (See STAR website for all tissues).

The collected datasets have versatile properties. Figure 2 depicts the number of locations, number of genes, sparsity level and number of datasets across 39 cohorts. As shown, there are substantial differences in the number of locations among cohorts analyzed by different techniques. Data from HDST has the largest number of locations, ranging from 118, 551 to 181, 367, which is consistent with the fact that HDST resolves 2*μm* spots [12] while ST resolves 100μm spots [11], and 10x Visium resolves 55 *μm* spots [27]. For imaging-based SRT technologies, seqFISH and seqFISH+ share a similar number of spots, while MERFISH can profile more cells. For the number of genes (Figure 2 (b)), seqFISH and MERFISH resolve 161 and 249 genes, respectively. SeqFISH+ resolves 10, 000 genes due to its implementation of 10, 000 gene probe sets. For sequencing-based SRT approaches, ST (including ST from 10x Visium) and HDST have a similar number of genes. ST from 10x Visium uses three gene probe sets, which include 32, 285, 36, 601 and 17, 943 genes, respectively. For sparsity (Figure 2 (c)), datasets resolved by ST and ST from 10x Visium have a high sparsity level from 59.96% to 94.72%, while datasets resolved by HDST have an even higher zero proportion from 99.96% to 99.99%. Datasets resolved by MERFISH and seqFISH+ have moderate sparsity levels, from 59.43% to 93.85%. seqFISH profiles datasets with low sparsity levels from 1.13% to 17.73%. For the number of datasets (Figure 2 (d)), while most of the 10x Visium datasets have only one dataset, mouse hypothalamus (n=36) and mouse hippocampus (n=21) have the largest number of datasets per cohort. Interestingly, when we visualize the number of genes and the number of locations for the collected datasets (all seven sequencing-based and imaging-based SRT techniques) chronologically from 2016 to 2022 (Figure S1), we notice a clear rising trend in the number of locations, which corresponds with the technological progress of increasingly high resolutions.

**Fig. 2.**
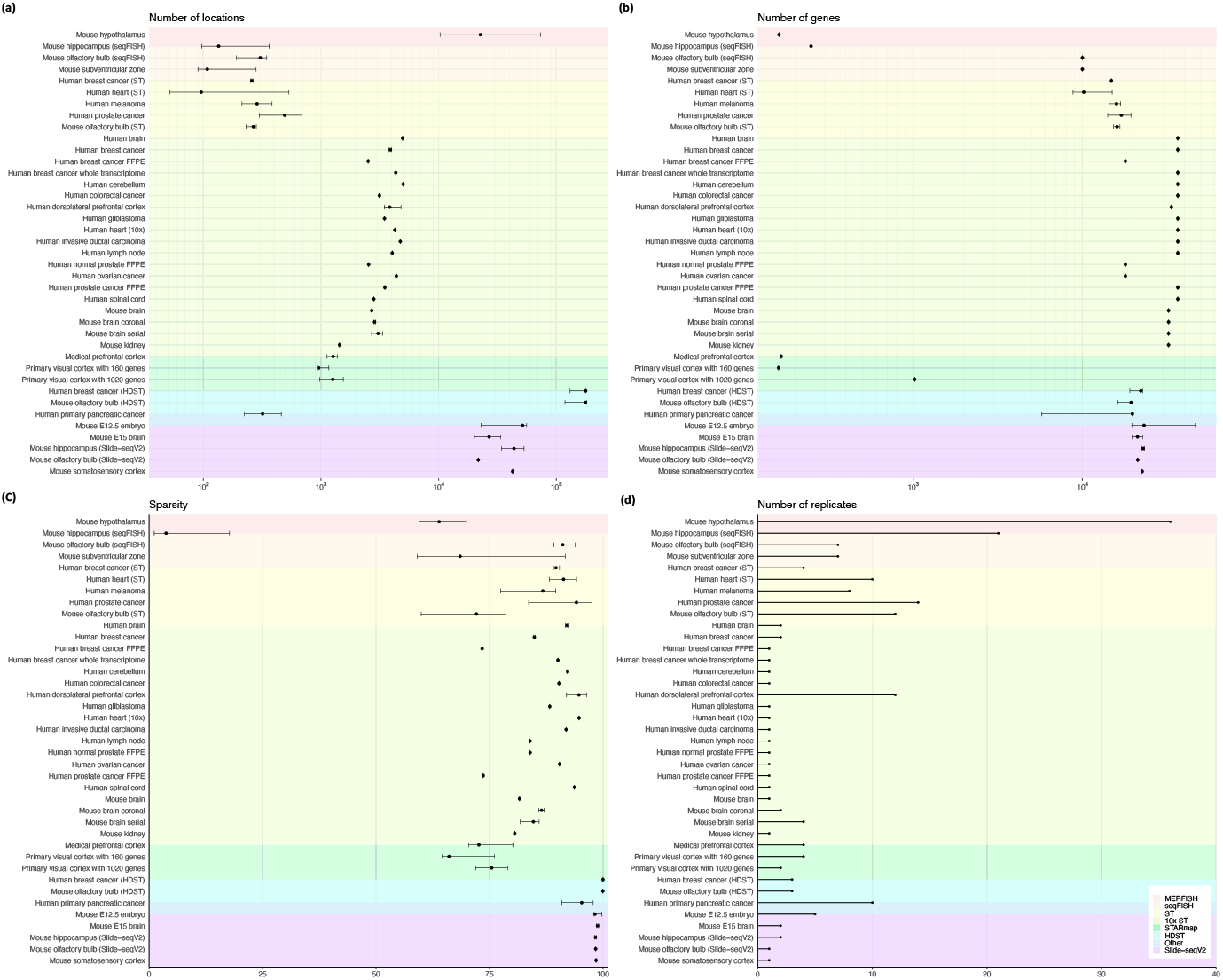
Scatter plots of the (a) number of locations (spots or cells), (b) number of genes, (c) sparsity level, and (d) number of datasets for 39 cohorts. The *y* axis on each plot shows the cohort names. The *x* axis depicts the size of log 10 transformed data property in (a) and (b) and raw quantity in (c) and (d). Scatter plots (a) to (c) point indicate the median of the corresponding data property with the range of that property within each cohort as an error bar. Background color depicts the SRT technologies that resolve the cohorts.

### 2.3 Benchmarking SV Gene Detection Methods

A key component of STAr is the unbiased assessment of the performance of SV gene detection methods. We utilized comprehensive metrics to evaluate several computational approaches that identify SV genes on an extensive range of simulation datasets. We have publicized this comprehensive collection of simulation data (see ‘Methods’), generated by five spatial patterns (spot, linear, MOB I, MOB II, and BC patterns, in Figure 3) and four zero-inflation settings (0, 10%, 30% and 50%). For each combination of pattern and zero-inflation settings, there are 30 replicated datasets. This sums up to 600 simulation datasets in all for benchmarking.

**Fig. 3.**
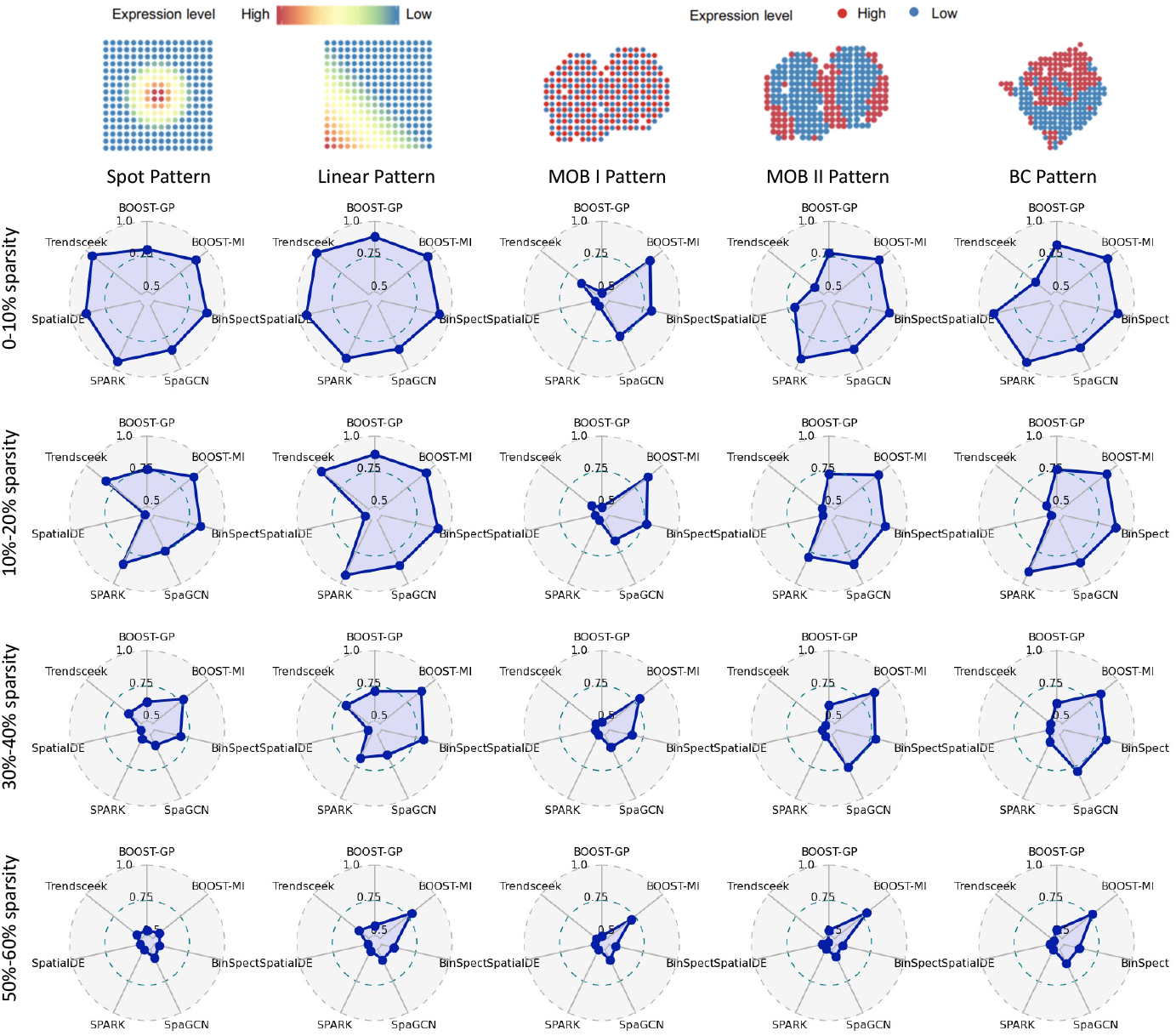
Radar plots of AUCs achieved by BOOST-GP, BOOST-MI, BinSpect, SpaGCN, SPARK, SpatialDE, and Trendsceek under 20 combinatorial scenarios from five spatial pattern and four sparsity settings.

We began by curating a comprehensive list of SV gene detection methods and then applied them to all of the simulation data. These methods were BOOST-GP [20], BOOST-MI [22], BinSpect [23], SpaGCN [24], SPARK [19], SpatialDE [18] and Trendsceek [21]. We summarized these methods (Table S2) and provided details of their implementation (Section S1 in Supplementary Notes) to ensure reproducibility. When a method had multiple running options, we selected the optimal settings. For instance, we chose mark-variogram as the summary statistic for Trendsceek and rank as the binarization method for BinSpect since those settings achieved the best performance for their methods respectively.

We then visualized the average area under the receiver operating characteristic curve (AUC) by different methods on 30 replicated datasets under each scenario (Figure 3). BOOST-GP, BOOST-MI and BinSpect achieved the best performance on average. SPARK has very high accuracy in non- and low-zero inflation but suffers from reduced power with increased sparsity. Compared to the other methods, SpatialDE and Trendsceek have unsatisfactory performance on the simulation data. SpaGCN performs better than SpatialDE and Trendsceek but is inferior compared to the other four methods. In addition, SV genes with the MOB I pattern can only be detected by BOOST-MI and BinSpect. Results evaluated by the *F*_1_ scores and Matthews correlation coefficients (MCC) are also available (Supplementary Notes Figure S2). For *F*_1_ scores and MCCs, each method requires a threshold to determine whether a gene is an SV gene or not. We set the selection criteria as adjusted p-values less than 0.05 for BinSpect, SpaGCN, SPARK, SpatialDE, and Trendsceek, and a Bayes factor greater than 150 for BOOST-GP and BOOST-MI. Similar to the AUC results, BOOST-GP, BOOST-MI and BinSpect achieved very stable performance across all settings. The computational cost of each method is listed in Table 1. The two Bayesian methods have a higher computational burden since they need multiple iterations to perform posterior sampling for the model parameters.

**Table 1:** Computational cost for BOOST-GP, BOOST-MI, BinSpect, SpaGCN, SPARK, SpatialDE and Trendsceek on simulated data.

| Methods | Range of running time per gene (in seconds) |
| --- | --- |
| BOOST-GP | 12.001 – 34.935 |
| BOOST-MI | 4.299 – 10.650 |
| BinSpect | 0.007 – 0.010 |
| SpaGCN | 0.012 – 0.014 |
| SPARK | 0.096 – 0.107 |
| SpatialDE | 0.035 – 0.052 |
| Trendsceek | 0.363 – 1.166 |

Given the general availability of the seven SV detection methods, we further explored two rank aggregation methods – GEO and MC2 [28] on all simulated data. Similarly, we calculated the averaged AUC, *F*_1_ score, and MCC (Supplementary Notes Figure S3). In general, although the two rank aggregation methods achieved similar performance, they are more robust under all settings except for the MOB I pattern.

Lastly, a detailed comparison visualized through bar charts is available online. This includes six performance metrics (See ‘Methods’ for details) and the computational cost.

### 2.4 Interactive SV Analysis Results

In addition to providing public SRT datasets and SV methods, STAr also displays collected results for all popular SV gene identification methods on 50 ST datasets across 21 cohorts, including the mouse olfactory bulb (12 datasets), human breast cancer (four datasets), human heart (ten datasets), and 24 other datasets from the 10x Visium platform. Users can select a cohort and a dataset, and a table for SV gene measurements will be shown. We provide the results from six SV gene identification methods: BOOST-GP, BOOST-MI, BinSpect, SpaGCN, SPARK and SpatialDE. Due to Trendsceek’s unfavorable performance on simulated data, we did not apply it to real datasets. For BOOST-GP and BOOST-MI, Bayes factors are displayed, while adjusted p-values are included in the table for the other four methods. Additionally, the result table includes the rank of SV genes from two rank aggregation methods: GEO and MC2. The p-values obtained from SpaGCN are excluded from GEO and MC2. SV genes in the union of 1, 000 top genes for five methods (excluding SpaGCN) were assigned a combined rank with ties. Genes that were not in the union of 1, 000 top genes, but had at least one result across the five methods, were ranked as the current maximum rank plus 1. The analysis results for all datasets are summarized in Table S4 in the Supplementary Notes. On the website, users can click the column title to sort the table in ascending or descending order by the corresponding column.

Moreover, STAr can also visualize spatial gene expression counts for these data. Users can select a gene of interest and the spatial expression counts will be shown on their associated hematoxylin and eosin (H&E)-stained tissue slide image. Points are located based on the location information from ST data and the color of the point represents the gene expression level. Users can modify the point symbol size and choose raw or normalized counts (normalization procedures in supplementary notes Section S2) to display; see Figure 4 for examples. Hovering over the point will display a pop-up box with the coordinates and expression value. A histogram next to the expression display figure shows the distribution of expression levels for the corresponding gene. Selecting a point in the expression display figure will highlight the corresponding bar in the histogram to show the position of expression level on this spot in the distribution of expression levels of all spots. These functions enable the user to conveniently explore the spatial characteristics of each gene from various ST data.

**Fig. 4.**
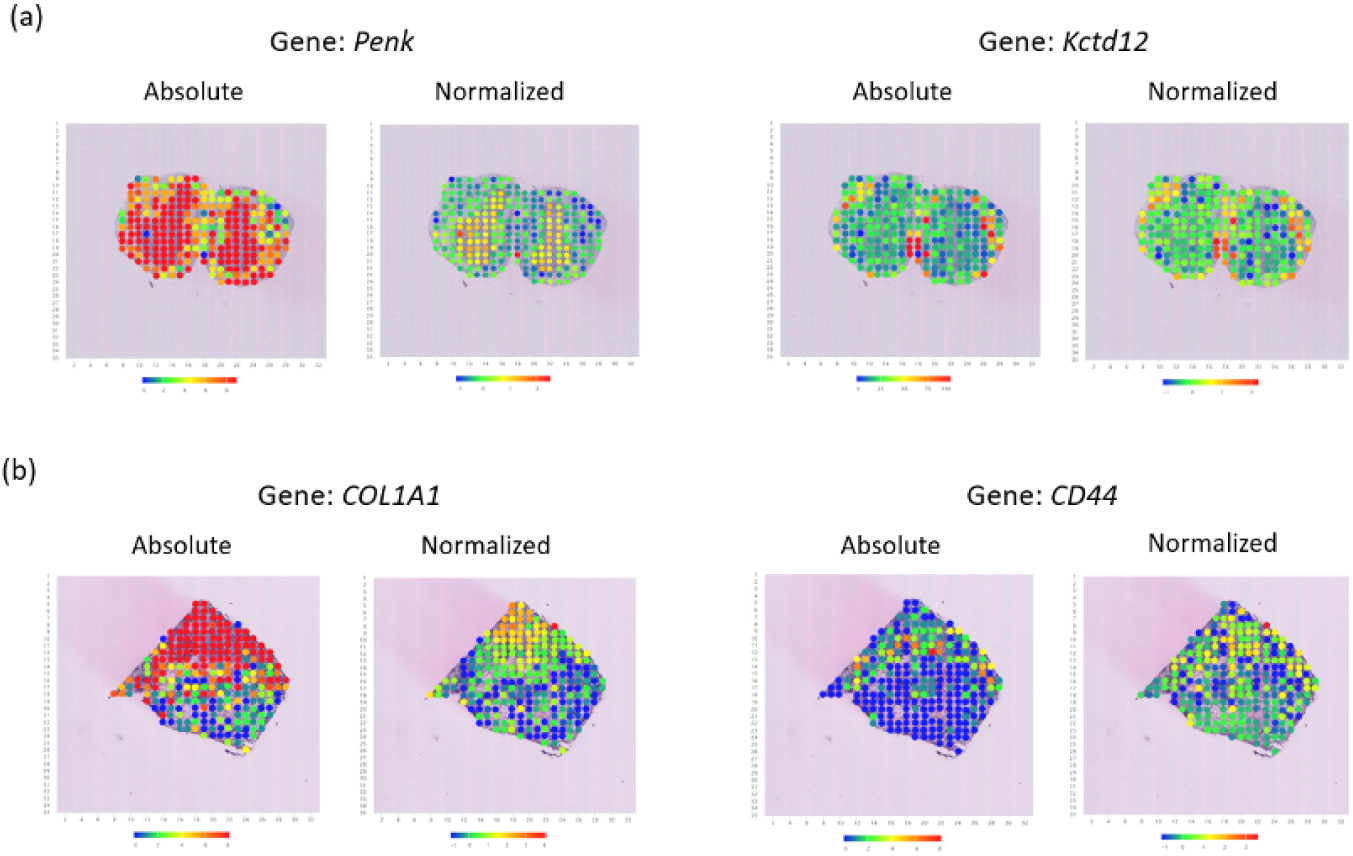
Examples of visual examination of gene expressions on STAr: Spatial expression pattern visualization on raw and normalized count data for four spatially variable genes identified by most of methods in (a) mouse olfactory bulb ST data and (b) human breast cancer ST data. Color represents gene expression levels (red, high; blue, low).

### 2.5 An Open Platform for User Contribution

STAr is intended as a dynamic resource. In its current stage, we provide three core components (Figure 5): 1) We curated datasets from publicly accessible resources or our in-house simulations; 2) Similarly, we surveyed SV gene analysis methods and incorporated them in a documented companion `R` package `boost` (https://boost-r.readthedocs.io/en/latest/). It provides a unified analysis routine that includes all benchmarked spatial gene analysis methods; 3) We applied all methods to curated datasets and released the analysis results. We ensured that the datasets, methods, and results can be viewed in STAr and downloaded without restriction. We also provide data descriptions in a tabular format as well as a spatial gene viewer.

**Fig. 5.**
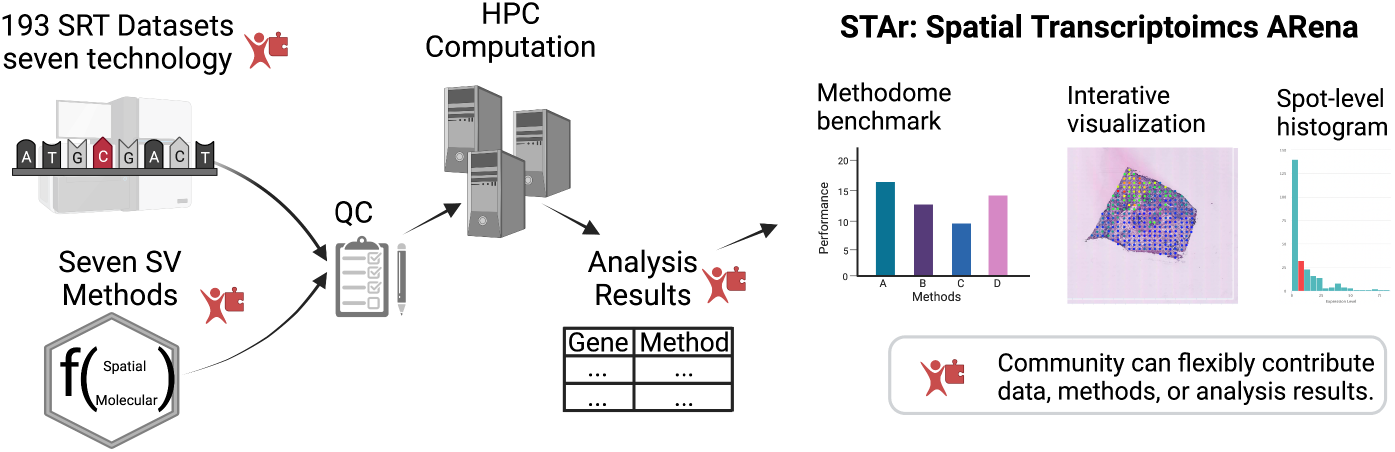
User contribution guidance: STAr allows users to upload data, methods, and analysis results to its main web portal. All contributed resources will need to provide metadata and undergo quality control. STAr will then perform reproducible analysis for all combinations of data and methods and the results will be displayed on the main website.

STAr provides interactive features to visually examine analysis results, offering an online viewer that supports the visualization of thousands of spots, optionally color-coded by specific gene expression amounts. Users may select a specific spatial location or spot to highlight the corresponding bars on the histogram. This real-time data depiction gives an overarching view of the spatial dynamics of a tissue’s transcriptome. To ensure a smooth online user experience, we designed the website pages and data to utilize static file contents on the server. Altogether, STAr offers a user-friendly website to browse SRT resources.

STAr can grow with the development of SRT technology. It welcomes contributions from communities, with clear contribution guidelines. New spatial transcriptomics data or methods can be submitted as long as appropriate metadata is provided as follows. For new datasets, we require the technology name, tissue location, disease type, and citation. For new method submissions, we require open-source code, a permissive license, and a software version. For analysis results, we require a documented dataset and method as described above. By reducing upload restrictions, the contributors can provide up-to-date research results, which can accelerate SRT research. We will update these resources periodically on the STAr website to fulfill our goal of maintaining a high-quality resource for basic biology and methodology research.

## 3 Methods

### 3.1 Real SRT Dataset Curation

We curated 193 real datasets in 39 cohorts from seven mainstream SRT technologies: ST, seqFISH/seqFISH+, MERFISH, Slide-seqV2, STARmap, HDST, and an integration of microarray and single-cell RNA sequencing (Other). A detailed summary can be found in Table S1 in the Supplementary Notes. Because the SRT data were generated by different labs and stored in different formats, we needed to process the raw data into the same data structure. Here, we have focused on the most available profiles among all the existing SRT datasets, including the molecular profiles (i.e., gene expression count table), geospatial profiles (i.e., coordinate information of spots or cells), and imaging profiles (H&E-stained Whole Slide Image) if available. Each cohort was further split into different datasets by individuals, tissue, section layers, or field of view.

The raw molecular and geospatial profiles were in .tsv, .txt, .xlsx, .csv, or .mtx formats. We downloaded the data, reorganized each molecular profile into an *n*-by-*p* count matrix *Y* and each geospatial profile into an n-by-2 numeric matrix *T*, and stored them in a .zip file to enable the coexistence of multi-type objects. Particularly, in each dataset, four objects were included as .csv formatted processed files: count matrix *Y* = [*y_ij_*]_*n×p*_, location matrix *T* = [*t_j_*]_*n*×2_, gene name, and gene ID. At a greater level of detail, the methods used to process the raw datasets for datasets resolved by different technologies were slightly different. The datasets resolved by HDST are extremely large with the numbers of spots over 170, 000, disallowing normal matrix transformation. Consequently, the count matrix of HDST datasets contains only non-zero entries in the .csv file. The datasets from 10x Visium were aligned based on barcode information because raw data from the 10x Visium website contain separate files including barcode information, count matrix and tissue positions. As for the mouse hypothalamus cohort data, 36 individual mice were processed separately. For cohorts resolved by seqFISH and seqFISH+, datasets were separated by fields of view, processed, and stored in the unified format.

Before analyzing a given spatially resolved count dataset, we first implemented a two-step quality control procedure. The first step was sample-wise quality control. In count data analysis, if the total number of reads for a cell or spot falls above or below specific values (discussed below), then this may indicate poor sequence quality owing to duplicate reads or limited sampling bias. For instance, let 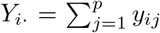 denote the total number of reads observed in sample *i*. A sample *i* will be removed if *y_i_* < *Q_i_* – 3(*Q*_3_ – *Q*_1_) or *y_i_* > *Q*_3_ + 3(*Q*_3_ – *Q*_1_), where *Q*_1_ and *Q*_3_ are the lower and upper quartiles (i.e., the 25th and 75th percentiles) of the total reads of all the cells or array spots (i.e., *y*_1_,..., *y_n_*). Note that in the context of box-and-whisker plots, a data point is defined as an extreme outlier if it falls outside these two limits. Quality control methods used in previous papers are as follows: For ST datasets, SPARK [19] filtered out spots with total read counts less than ten, while Trendsceek[21] excluded spots with less than five read counts. For the MERFISH dataset, SPARK removed ambiguous cells that were identified as putative doublets. For the seqFISH dataset, SPARK and SpatialDE [18] filtered out cells with *x*-or *y*-axis values falling outside the range of 203 — 822 pixels to eliminate edge artifacts. The second step was feature-wise quality control. A common procedure in SRT dataset analysis is to filter out genes of extremely low abundance. For instance, SPARK excluded genes expressed in less than 10% of the array spots. Similarly, SpatialDE filtered out genes with expression in less than three array spots.

### 3.2 Simulated Dataset Generative Model

We followed the same data generation schemes described in the SPARK [19], BOOST-GP [20] and BOOST-MI [22] papers. The benchmark simulated data was generated from three artificial spatial patterns and two real spatial patterns, which are depicted in Figure 3. The first two artificial patterns, named spot and linear, were on a 16 × 16 square lattice (*n* = 256 spots), while the remaining one named MOB I was on the *n* = 260 spots in the mouse olfactory bulb study (MOB). The MOB II and BC patterns were on the *n* = 260 and 250 spots, respectively. They were respectively constructed from the MOB study and human breast cancer (BC) study. In total, there were 100 genes, among which 15 were spatially variable (SV) genes.

In the following, we denote the gene expression count table as an *n* × *p* matrix *Y*. Each entry 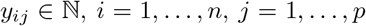 is the read count for gene *j* observed at spot *i*. We also denote the underlying normalized expression levels as a *n* × *p* matrix Λ. For each gene *j*, the latent normalized expression level at spot *i* λ_*ij*_ was generated via,

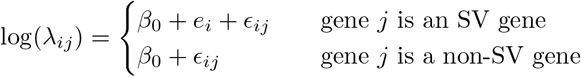

where *β*_0_ is the baseline of log-normalized expression levels and *ϵ_ij_* is the non-spatial errors following *ϵ_ij_* ~ *N*(0, *σ*^2^). We set *β*_0_ = 2 and *σ* = 0.3. Thus, for a non-SV gene, the normalized expression levels were independently and identically distributed (i.i.d.) from a log-normal (LN) distribution with a mean and variance of 2 and 0.3^2^ , respectively. Consequently, no spatial correlation should be observed. For the spot pattern, the values of spatial pattern effect *e_i_*’s of the four center spots, i.e., (8, 8), (8, 9), (9, 8), and (9, 9), was set to log 6, while all others were linearly decreased to 0 within the radius of five spots. For the linear pattern, the value of *e_i_*’s of the most bottom-left spot, i.e., (1,1), was set to log6, while all others were linearly decreased to 0 along the diagonal line. For the remaining three patterns, each spot was dichotomized into one of two groups: high and low expression levels with *e_i_* = log 3 and 0, respectively. Note that the artificially generated MOB I pattern exhibits a complete attraction spatial pattern (the clustering of points with different expression levels), while the MOB II and BC patterns display the opposite spatial structure. To characterize the excess zeros and over-dispersion in the sequencing count data, we sampled the observed gene expression counts *y_ij_*’s from zero-inflated negative binomial (ZINB) distributions,

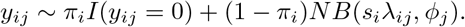

The size factors *s_i_*’s were i.i.d. from *LN*(0,0.2^2^) and the dispersion parameters *ϕ_j_*’s are i.i.d. from Exp(0.1). For the choice of the false zero proportion *π_i_*, we randomly selected 0%, 10%, 30%, or 50% counts and forced their values to zero. Combined with the five patterns (i.e., spot, linear, MOB I, MOB II, and BC) and four zero-inflation settings, there were 5 × 4 = 20 different scenarios in total. For each of the scenarios, we independently repeated the above steps to generate 30 replicates.

### 3.3 SV Gene Detection Methods

SV gene identification is a critical aspect of downstream analysis for SRT techniques, providing a comprehensive exploration of SRT datasets and contributing to new discoveries for further genomics research. To evaluate and exhibit cutting-edge statistical methods for spatially variable (SV) gene analysis in an unbiased manner, we collected seven existing statistical methods proposed for identifying SV genes and classified them into three main categories according to how they define the spatial correlation of gene expressions when modeling the SRT data.

1) Kernel-based spatial models: Model the gene expression levels via the Gaussian process and quantify the spatial dependency via various kernel functions. SpatialDE [18] is a kernel-based method, modeling the normalized expressions with a Gaussian process to identify SV genes. It applies different spatial kernels, including the squared exponential, periodic, and linear, to capture spatial dependency. SPARK [19] is another kernel-based approach. However, it directly models the gene expression counts via a generalized linear model and employs spatial kernels with different length-scale characteristics to accommodate various spatial patterns. It has higher power compared to SpatialDE. Moreover, both approaches have low computational cost. BOOST-GP [20] is a Bayesian hierarchical model also via the Gaussian process to identify SV genes. Unlike SPARK, it models gene expression counts with a zero-inflated negative binomial distribution, so it achieves better performance on highly sparse SRT data. However, the computational cost of BOOST-GP is much higher since it is under the Bayesian framework. For all three above approaches based on Gaussian processes, kernel functions need to be pre-defined, which does not address the task of discovering spatial patterns that cannot be defined by kernel functions.

2) Energy-based spatial models: BOOST-MI [22] is a multi-stage analysis tool that identifies SV genes by quantifying spatial correlation via the energy measurement in the Ising model. It dichotomizes relative expression levels and characterizes the resulting gene-specific binary spatial pattern via inferring the Ising model interaction parameter under a Bayesian framework. BOOST-MI provides more robust detection results and can detect more potential spatial patterns of gene expressions, especially for those patterns that cannot be defined by kernel functions. It can also spontaneously overcome the high sparsity issue in real SRT data. However, BOOST-MI is only feasible on SRT data obtained from sequencing-based techniques, whose gene expression levels are measured approximately on a lattice grid.

3) Other spatial analysis methods: There also exist other approaches proposed for SV gene detection. Trendsceek [21] is a nonparametric method without distribution specification, which models data via a marked point process. SV genes are identified by mark-segregation hypothesis testing and testing is carried out with permutation tests. Trendsceek is computationally intensive and has noticeably unsatisfactory performance compared to other model-based methods [19]. BinSpect [23] is a non-model-based approach based on the statistical enrichment of spatial network neighbors. The framework of BinSpect is straightforward and it has the highest computational efficiency compared to the other five methods described above. However, as mentioned in the BOOST-MI paper [22], it is an aggressive method that may suffer some misclassification bias. SpaGCN [24] is a spatial domain and spatially variable (SV) gene identification approach based on a graph convolutional network. Unlike other SV gene detection methods, it first integrates information from the gene expression profile, location, and pathology image to identify spatial domains. Then, SV genes are defined based on domain-guided differential expression analysis.

### 3.4 Rank Aggregation Methods

Rank aggregation is the process of combining multiple ranking lists (known as “base ranker”) into a single ranking list (known as “aggregated ranker”). In recent years, rank aggregation methods have emerged as an important tool for integrating information from individual genomics studies that address the same biological question. We applied two rank aggregation methods, GEO and MC2, to combine SV gene identification results from methods in both simulated and real data.

GEO is one of the rank aggregation methods in Borda’s method [29], which uses the geometric mean of base rankers to aggregate rank data.

MC2 is a rank aggregation method developed under a Markov chain modeling framework, where the union of items from all base rankers forms the state space. The transition matrix is then constructed in a way such that the chain will move to a state with better rankings in at least half of the base rankers and its stationary distribution will have larger probabilities for states that are ranked better. Hence, the aggregated rank is determined by the stationary probability of each state [29].

Figure S4 in the Supplementary Notes shows the AUCs achieved by several common rank aggregation methods under different scenarios from the simulated data. Robust rank aggregation (RRA) [30] is based on the comparison of actual ranked lists with a null model that assumes random order of input lists. A numerical score is assigned to each item based on the reference distributions of order statistics (i.e., beta distributions). The aggregated rank is obtained by sorting p-values. p-values are computed on the base of Bonferroni correction of the numerical scores to avoid the intensive computation required to obtain exact p-values. Stuart [31] identifies pairs of genes whose expression is significantly correlated in multiple organisms and then ranks them according to the Pearson correlation coefficient. We calculated p-values based on the distributions of order statistics. Summary statistics namely, mean, median, geometric mean, and L2-norm are denoted as MEAN, MED, GEO, and L2, where lower case letter ‘`r`’ for `R` package `RobustRankAggreg` and `t` for `R` package `TopKLists` since these two packages have different implementations for MEAN, MED, and GEO. MC2 was implemented in `TopKLists` package in `R`. From Figure S4 in the Supplementary Notes, tGEO and MC2 have relatively better performance than others, which led us to choose these two rank aggregation methods for displaying the aggregated rank on our website.

### 3.5 Performance Metrics

We used various metrics to evaluate the accuracy and efficiency of different SV gene identification methods on the simulated data, for which the ground truth is known. As the outcome is binary (i.e., whether a gene is an SV gene or not), we considered six widely used performance metrics for binary classifiers: the area under the curve (AUC) of the receiver operating characteristics (ROC), sensitivity, specificity, false discovery rate (FDR), F_1_ score, and the Matthews correlation coefficient (MCC) [32]. The AUC considers both true positive (TP) and false positive (FP) rates across various threshold settings, while the remaining balance TP, FP, true negative (TN), and false negative (FN), pinpointing a threshold. In particular, we chose a significance level of 0.05 to select SV genes based on adjusted p-values for BinSpect, SPARK, SpatialDE, and Trendsceek, while we set a Bayes factor cutoff of 150 for the two Bayesian methods, BOOST-GP and BOOST-MI. In the context of the SV gene identification problem, TP, FP, TN, and FN are defined as the number of true SV genes that are identified, the number of true non-SV genes that are identified, the number of true SV genes that are not identified, and the number of true non-SV genes that are identified, respectively. We briefly introduce each performance metric in Table 2.

**Table 2.** Performance metrics for SV gene identification approaches evaluation.

| Formula | Description | Range |
| --- | --- | --- |
| $\text{sensitivity} = \frac{TP}{TP+FN}$ | Proportion of correctly identified SV genes across all true SV genes | 0 to 1 |
| $\text{specificity} = \frac{TN}{TN+FP}$ | Proportion of correctly identified non-SV genes across all true non-SV genes | 0 to 1 |
| $FDR = \frac{FP}{TP+FP}$ | Proportion of wrongly identified SV genes across all identified SV genes | 0 to 1 |
| $F_1 = 2 \times \frac{TP}{2 \times TP + FP + FN}$ | Harmonic mean of the precision and sensitivity | 0 to 1 |
| $MCC = \frac{TP \times TN - FP \times FN}{\sqrt{(TP+FP)(TP+FN)(TN+FP)(TN+FN)}}$ | Correlation coefficient between the identified and true SV gene classifications | -1 to 1 |
| $AUC = \int_{x=0}^1 TPR(FPR^{-1}(x))dx$ | Measure of performance across all possible classification thresholds | 0 to 1 |

Notes: $TPR=TP/(TP+FN)$ ; $FPR=FP/(FP+TN)$

For the efficiency evaluation, we ran each method with its default setting on a high-performance computing server with two Intel Xeon CPUs (45 MB cache and 2.10 GHz) and 250 GB memory and recorded its runtime in seconds.

Note that BOOST-GP, BOOST-MI, SPARK, BinSpect, and Trendsceek are written in `R`, the first three of which use `Rcpp` to accelerate computation, while SpatialDE and SpaGCN are written in Python.

## 4 Discussion and Conclusion

SRT is prevailing across many biomedical fields, offering a novel perspective to quantitative biological research. Suitable statistical analysis is critically needed for biological discoveries to efficiently utilize these technological advancements. We have observed an increased interest in SRT data generation or curation (e.g., SpatialDB [33]), as well as methodological development for various biological questions (e.g., tissue spatial domain detection [34], gene spatial expression patterns identification [18, 19], cell-type decomposition [35, 36]. We offer STAr, a unique resource that provides curated SRT datasets (including real and simulated datasets), widely used SV gene analysis methods, and SV analysis results to the community.

In this study, we have created an open-source and general platform to facilitate spatially resolved transcriptomics research. Our contribution includes a large collection of curated spatial transcriptomics datasets drawing from 39 cohorts in total (193 replicates, layers, samples or sections). These are from 16 tissues (ten organs) from humans or mice, spanning from health controls to eight disease subtypes. We provide both the original and processed datasets, where a quality control procedure was applied to the latter. Additionally, we also provide 600 simulated data replicates under 20 simulation settings with five spatial gene patterns and four sparsity settings where the true SV genes were known *a priori*. Based on these data resources, we extensively benchmarked spatial gene detection methods using different statistical measurements. To facilitate a benchmark, we provide the analysis results in tabular form with extensive analysis results, as well as a spatial gene visualizer, with interactive image-based spatial gene expression. All resources (i.e., datasets, methods, and analysis results) are publicly available.

Most importantly, our platform is continuously evolving. We welcome contributions from users in terms of new data, methods, and analysis results. In our contribution guidelines, we specify that these submissions are of adequate research quality (e.g., reproducibility). This growing list of resources can further expand research in the spatial transcriptomics community. We envision that STAr will curate a diverse set of simulation and real datasets, detail state-of-the-art statistical methods, and provide an unbiased evaluation of these methods. As a result, STAr will help accelerate future SRT methodological innovations that aim to advance quantitative biomedical science.

## Availability of Data and Materials

The lists of public SRT datasets are collected from from SciLifeLab (https://www.scilifelab.se/data), 10x Genomics (https://www.10xgenomics.com/products/spatial-gene-expression), the Broad Institute of MIT and Harvard (https://www.broadinstitute.org/), Dryad (https://datadryad.org), and GitHub repositories. All real and simulated SRT datasets can be downloaded from STAr website.

Our SV gene analysis pipeline is available via an `R` package `boost` published on GitHub (https://github.com/estfernan/boost).

## Supplementary Materials

**Figure S1**. The boxplots of AUCs achieved by rank aggregation methods. **Figure S2**. Scatter plots of number of genes and number of locations for datasets from 2016 to 2022. **Figure S3**. Radar plots of *F*_1_ scores MCCs achieved by seven spatially variable gene identification methods. **Figure S4**. Radar plots of AUCs, Fi scores and MCCs achieved by rank aggregation methods. **Table S1**. Summary of collected real SRT data. **Table S2**. Summary of spatially variable gene analysis methods. **Table S3**. Performance metrics for seven spatial gene detection methods and two rank aggregation methods. **Table S4**. Analysis results for 50 real datasets. **Supplementary notes S1**. Implementation details of SV gene identification methods. **Supplementary notes S2**. Normalization procedure to obtain relative expression.

