## Supplementary materials for "Spatial Transcriptomics Arena (STAr): an Integrated Platform for Spatial Transcriptomics Methodology Research"

**Figure S1**. The boxplots of AUCs achieved by rank aggregation methods under the scenarios of two spatial patterns (BC and MOB II patterns in Figure 2).

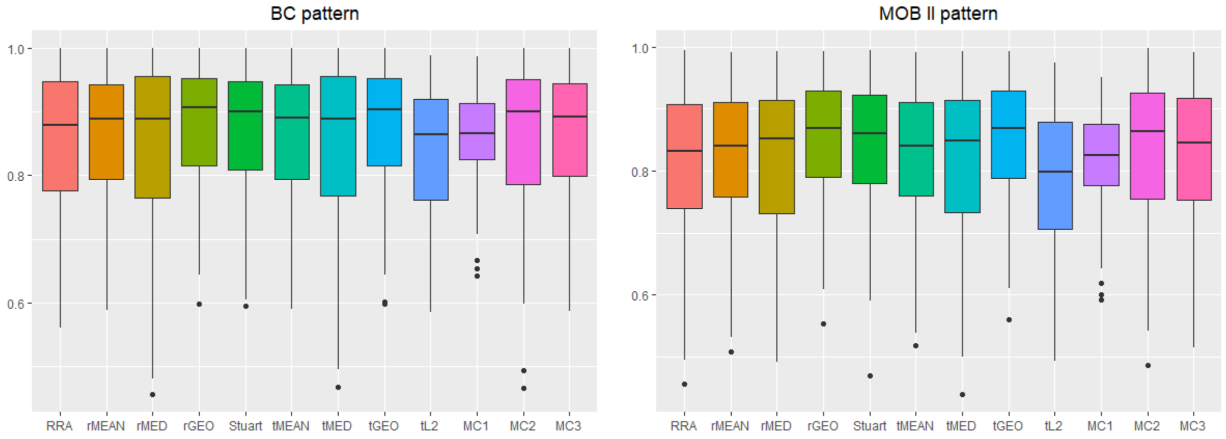

**Figure S2**. Scatter plots of number of genes and number of locations for datasets from 2016 to 2022: include seven sequencing- and imaging-based SRT techniques. Each color represents one technology.

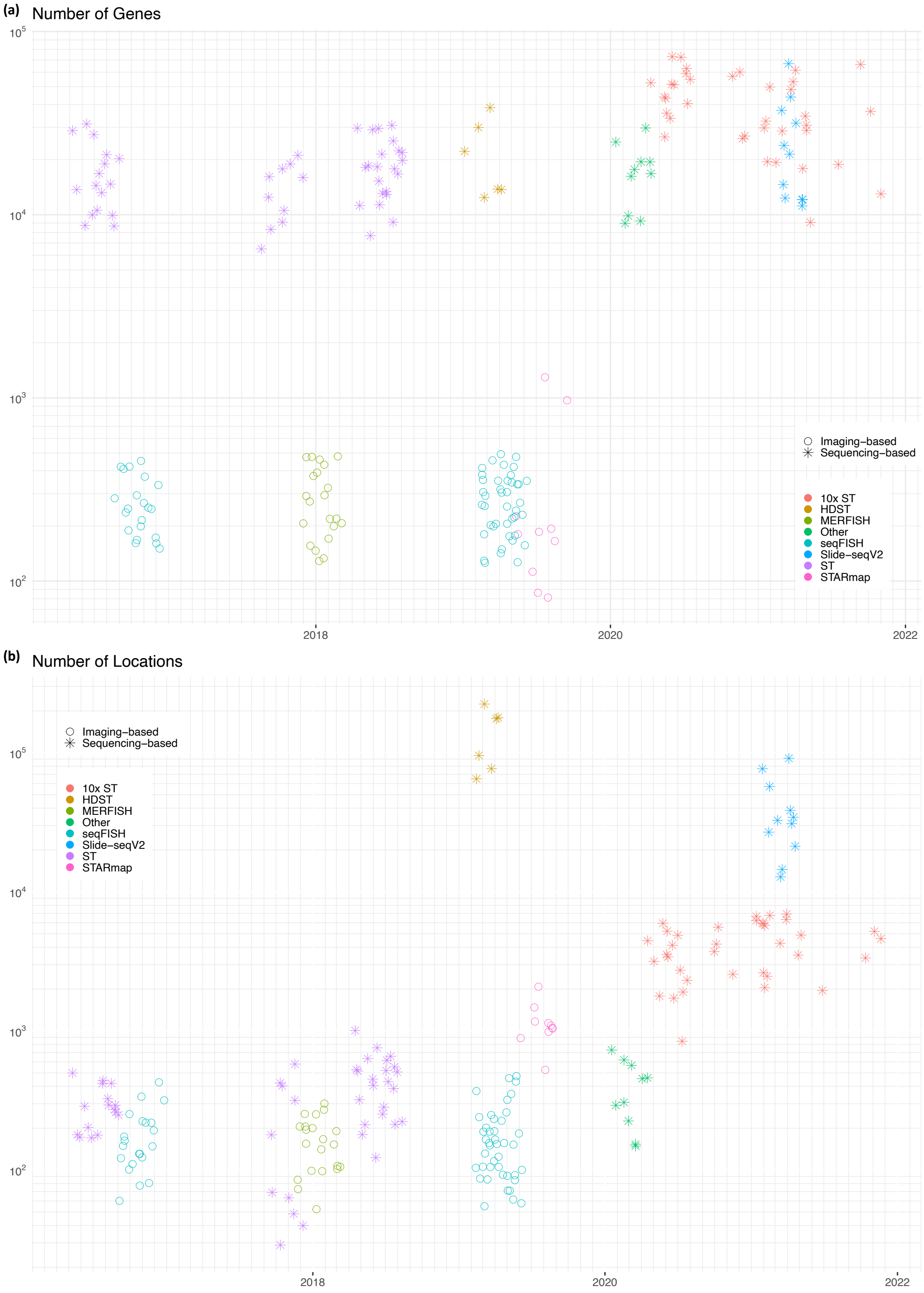

**Figure S3.** Radar plots of (a) $F_1$ scores and (b) MCCs achieved by seven spatially variable gene identification methods (BOOST-GP, BOOST-MI, BinSpect, SpaGCN, SPARK, SpatialDE, and Trendsceek) under different scenarios in terms of spatial pattern and zero-inflation setting.

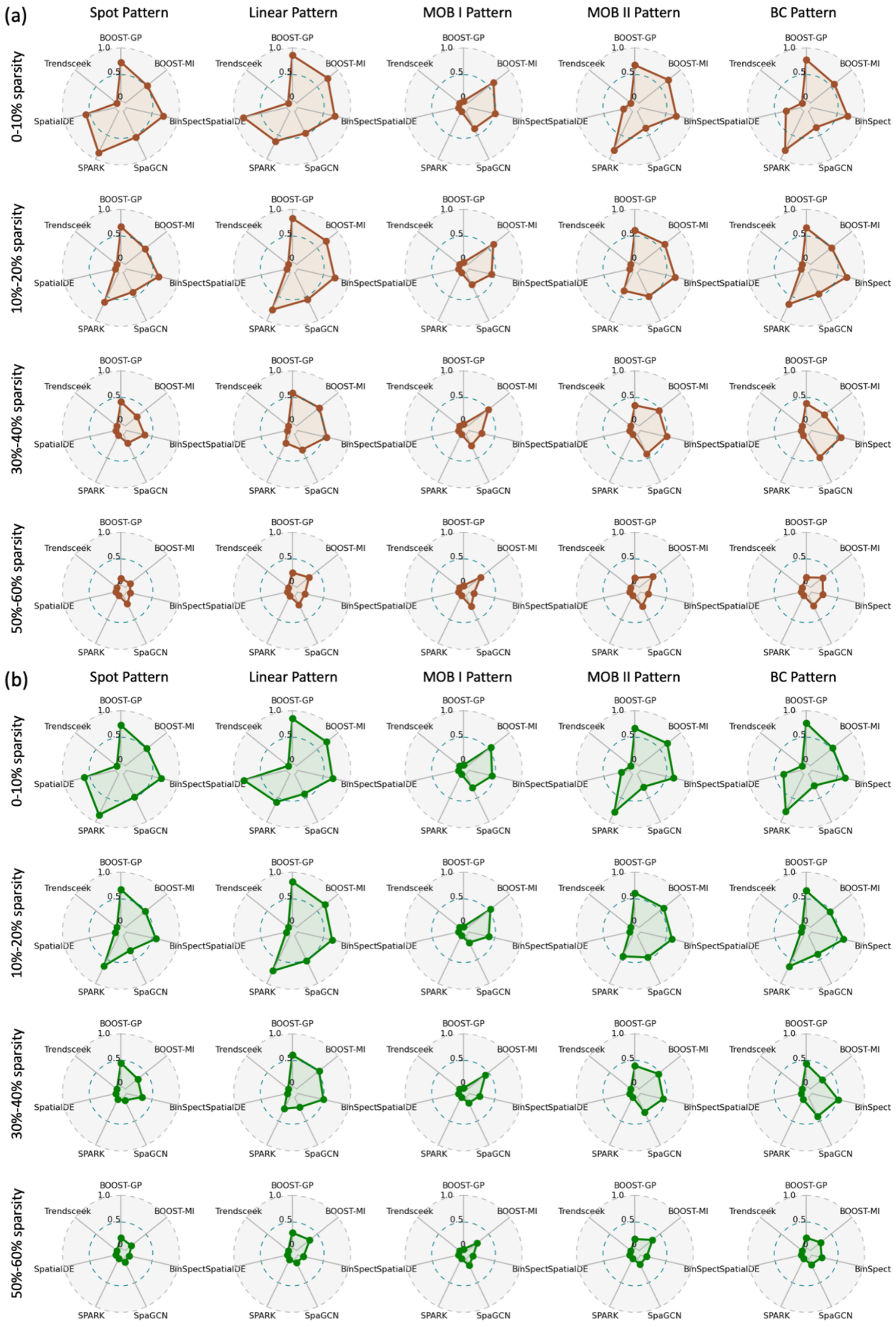

**Figure S4**. Radar plots of AUCs, $F_1$ scores and MCCs achieved by rank aggregation methods (a) GEO and (b) MC2 under different scenarios in terms of spatial pattern and zero-inflation setting.

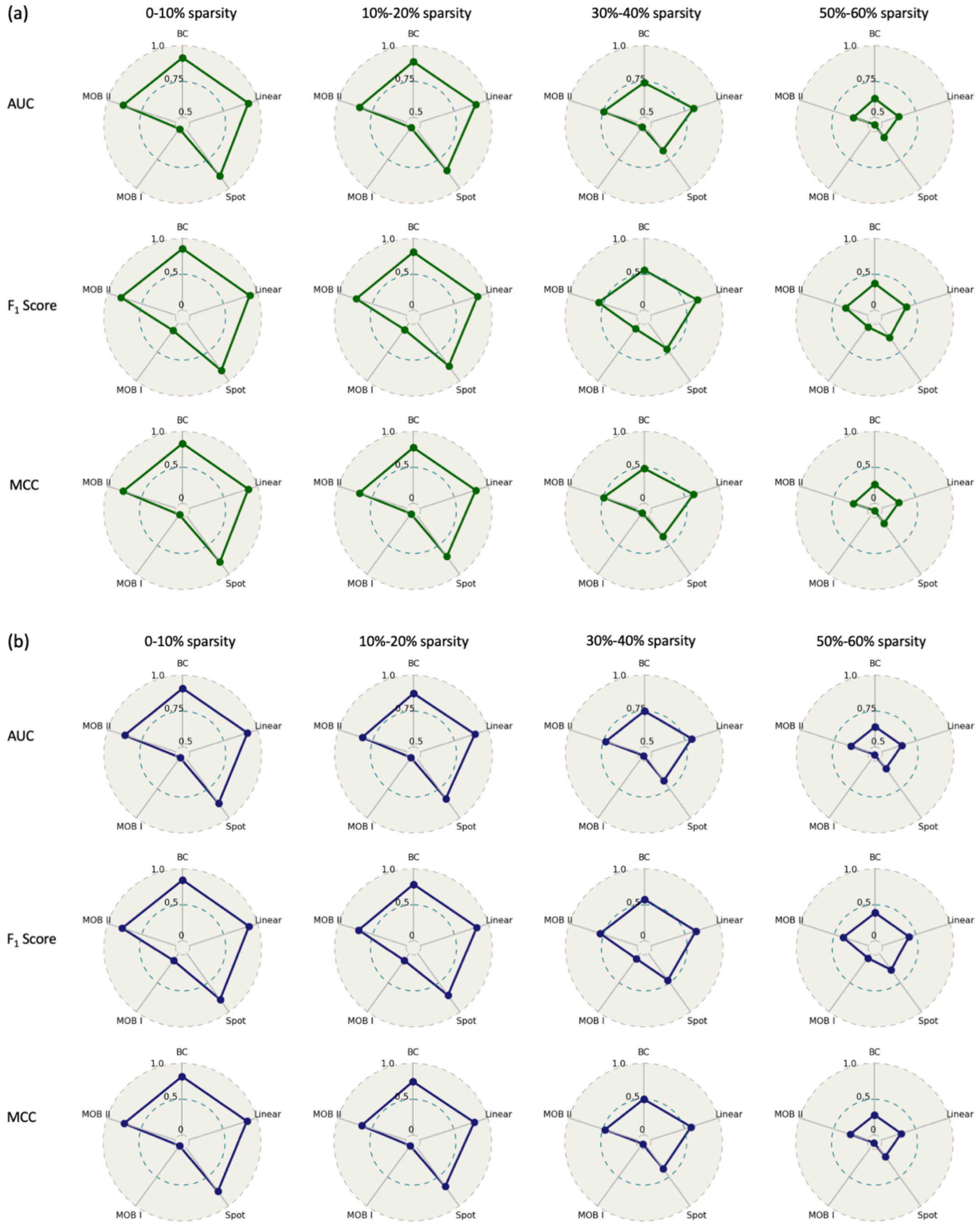

**Table S1.** Summary of collected real SRT data: summarizes the information from each cohort, including the source of publication, the technology used, profiling resolution, number of datasets, number of locations, number of genes, number of covariates, and sparsity level. Background color dissociates the technologies used to profile the cohorts.

supplementary_table_1.xlsx

**Table S2.** Summary of spatially variable gene analysis methods

| Name | Description | Version | Publication |
|---|---|---|---|
| BOOST-GP | A Bayesian hierarchical model where expression values are modeled as zero-inflated negative binomial distribution. | 1.0.0 | (Li et al., 2021) |
| BOOST-MI | A Bayesian hierarchical model where binarized gene expressions are investigated on a lattice grid. Genes with high interactions are reported. | 1.0.0 | (Jiang et al., 2021) |
| BinSpect | A multi-stage model to screen for genes that are enriched in binarized expressions in their neighbors. | 1.0.3 | (Dries et al., 2021) |
| SpaGCN | A graph convolutional network approach which integrates the information from gene expression profile, location, and pathology image to identify spatial domains. Genes with large expression fold changes between spatial domains are reported. | 1.2.0 | (Hu et al., 2021) |
| SPARK | A Gaussian process based hierarchical model where discrete expression values are modeled as Poisson distribution. Genes estimated with non-zero spatial kernel are reported. | 1.1.2 | (Sun et al., 2020) |
| SpatialDE | A Gaussian process based hierarchical model for normalized expression. Genes estimated with non-zero spatial kernel are reported. | 1.1.3 | (Svensson et al., 2018) |
| Trendsceek | A nonparametric method relies on marked point processes and is based on permutations to obtain p-values for non-parametric test statistics. | 1.0.0 | (Edsgärd et al., 2018) |

**Table S3.** Performance metrics for seven spatial gene detection methods and two rank aggregation methods: includes AUCs, $F_1$ scores, sensitivity, specificity, FDRs, MCCs, and run time for results from BOOST-GP, BOOST-MI, BinSpect, SpaGCN, SPARK, SpatialDE, Trendsceek, GEO, and MC2 under different scenarios in terms of spatial pattern and zero-inflation setting. The GEO and MC2 methods are rank aggregation methods based on the results of BOOST-GP, BOOST-MI, BinSpect, SPARK and SpatialDE.

supplementary_table_3.csv

**Table S4.** Analysis results for 50 real datasets: provides multi-sheet Excel file with spatially variable gene identification criteria for different methods. Criteria are illustrated by the reported p-values for BinSpect, SPARK, SpaGCN and SpatialDE, Bayesian factors for BOOST-GP and BOOST-MI, and ranks for two rank aggregation methods GEO and MC2.

supplementary_table_4.xlsx

### Supplementary Notes

#### S1. Implementation details of SV gene identification methods

We provide the details to implement methods for identifying SV genes.

We implemented BOOST-GP (Li et al., 2021) based on the codes provided on the GitHub repository (https://github.com/Minzhe/BOOST-GP). We input the sum of expression counts across all genes at each location as the size factor. All prior specifications of BOOST-GP were its default settings. To be specific, we set $a_\pi = b_\pi = 1$, $a_\phi = b_\phi = a_l = b_l = 0.001$, $a_\sigma = 3$, $b_\sigma = 1$, $a_\omega = 0.1$ and $b_\omega = 1.9$. For the posterior sampling, we set the default number of iterations as 1000, and first half of the iterations as burn-in. We applied the Bayes factor as the criteria of SV genes, which can be calculated from the function's output PPI (posterior probabilities of inclusion): Bayes factor $= \frac{PPI}{1-PPI} \times \frac{b_\omega}{a_\omega}$.

Boost-MI (Jiang et al., 2021) was run based on the codes provided on the GitHub repository (https://github.com/Xijiang1997/BOOST-MI). There were two pre-processing steps including 1) normalizing count data by applying the scale factor as the summation of total counts across all genes for sample (i.e., total sum scaling, or TSS); and 2) clustering the normalized expression levels into binary levels via the Gaussian mixture clustering (GMC). Then the modified Ising model was fit. The setting of priors of parameters was consistent with the suggestions in BOOST-MI paper. To be specific, the prior specifications were $\omega_0 \sim N(1, \tau_\omega^2)$ and $\theta \sim N\left(0, \tau_\theta^2\right)$ with $\tau_\omega = 2.5$ and $\tau_\theta = 1/3$. We set the number of iterations as 10000 and their first half as burn-in. The function outputs the Bayes factor for favoring the attraction pattern or repulsion pattern. For real datasets, we assigned the displayed Bayes factor for SV gene detection as the maximum of these two Bayes factors.

BinSpect (Dries et al., 2021) was implemented via the *R* package 'Giotto'. First, we obtained the normalized expression count matrix via the function 'normalizeGiotto', which is the Z-scoring of count matrix by genes and cells after log transformation of count matrix adjusted by scaling factor 6,000. Then, we applied ‘createSpatialNetwork’ to construct a network with minimum nearest neighbors set to two, and distance cut off for nearest neighbors to consider for Delaunay network set to 400. Then, ‘binSpect’ function was applied using either k-means or rank. Adjusted p-values were the criteria for the SV gene detection.

SpaGCN (Hu et al., 2021) was constructed under an open-source Python machine learning framework ‘PyTorch’. We implemented it via the Python package ‘SpaGCN’. For simulated data, the histology images were not available, so we set ‘histology = False’ when obtaining the adjacent matrix. For real data from 10x Visium, we also input their histology images. Before training the model, we filtered out spots with non-zero counts less than three and some special genes using functions ‘prefilter_genes’ and ‘prefilter_specialgenes’, respectively. Then we set its hyper-parameters using its default settings. Since we aimed at detecting global SV genes, we set the number of clusters to two for all simulated and real data. When training SpaGCN, we adjusted the maximum epochs to guarantee that the training process was converged. Then we obtained the refined spatial domains via functions ‘predict’ and ‘refine’. To obtain SV genes, we used functions ‘find_neighbor_clusters’ and ‘rank_genes_groups’ for two clusters and defined the SV genes as having differential expression on either of the spatial domain. The minimum adjusted p-values for two domains were the criteria to select the SV genes.

SPARK (Sun et al., 2020) was implemented through the R package ‘SPARK’. The original count matrix was filtered using the same procedure as described in the data quality control section by filtering out spots with total counts less than ten and filtering out genes which expressed in less than 10% array of spots. The input library size was calculated as the total expression counts of each spot across all filtered genes. Then we applied SPARK using its default settings. Ten kernel functions to model the spatial correlation were the same as described in its paper. We applied the adjusted combined p-values to select the SV genes.

We implemented SpatialDE (Svensson et al., 2018) through the Python package ‘SpatialDE’. Since it requires the normalized count matrix as the input, we transformed the gene expression counts to relative expressions using Anscombe’s variance-stabilizing transformation mentioned in the following section using functions in the 'NaiveDE' package. Then we run SpatialDE using its default setting. Q-values for multiple testing are the criteria for the SV gene detection.

We applied R package 'Trendsceek' to implement Trendsceek (Edsgärd et al., 2018) method. Before running the permutation tests, we pre-processed the data by the two steps: 1) We converted positions to point-pattern using function 'pos2pp'; and 2) We set the mark distribution of a point pattern via function 'set_marks'. Then we run Trendsceek using function 'trendsceek_test'. There are four sub-methods (conditional mean (E-mark), conditional variance (V-mark), Stoyan's mark correlation, and the mark-variogram) for Trendsceek. Results on simulated data showed that Trendsceek with mark-variogram has a significantly better performance than the other three sub-methods. Therefore, we set the output adjusted p-values from Trendsceek with mark-variogram as the criteria for SV gene selection.

#### S2. Normalization procedure to obtain relative expression

For sequencing count data, normalization is a necessary step for the development of SV gene analysis techniques to eliminate the influence of various sequence artifacts and biases. To obtain normally distributed and independent observations across cells, we performed data normalization as a data pre-processing step on our website. There are two steps for the normalization procedure. The first step entails transforming the count data, following a negative binomial distribution, into an approximately normal distribution using a variance-stabilizing transformation. Here we applied the approximate Anscombe's transform $\tilde{y}_{ij} = \ln(y_{ij} + \frac{1}{2\varphi})$, where φ is the overdispersion parameter of the negative binomial distribution and can be estimated by fitting the quadratic $s_j^2 = \bar{y}_j + \varphi\bar{y}_j^2$ across all genes (Anscombe, 1948; Svensson et al., 2018), where $s_j^2$ is the sample variance of gene j and $\bar{y}_j$ is the mean expression of gene j. Then, the relative gene expression levels are adjusted for the log-scale total read counts via a linear regression model to avoid the variation in cell size. For users' convenience of analysis, our website provides the normalized simulation data and the option of visualizing normalized real ST data.
